# Methotrexate inhibition of muropeptide transporter SLC46A2 controls psoriatic skin inflammation

**DOI:** 10.1101/2022.09.29.509906

**Authors:** Ravi Bharadwaj, Christina F. Lusi, Siavash Mashayekh, Abhinit Nagar, Malireddi Subbarao, Griffin I. Kane, Kimberly Wodzanowski, Ashley Brown, Kendi Okuda, Amanda Monahan, Donggi Paik, Anubhab Nandy, Madison Anonick, William E. Goldman, Thirumala-Devi Kanneganti, Megan H. Orzalli, Catherine Leimkuhler Grimes, Prabhani U. Atukorale, Neal Silverman

## Abstract

Cytosolic innate immune sensing is critical for protecting barrier tissues. NOD1 and NOD2 are cytosolic sensors of small peptidoglycan fragments (muropeptides) derived from the bacterial cell wall. These muropeptides enter cells, especially epithelial cells, through unclear mechanisms. We previously implicated SLC46 transporters in muropeptide transport in *Drosophila* immunity. Here we focus on *Slc46a2,* which is highly expressed in mammalian epidermal keratinocytes, and show that it is critical for delivery of DAP-muropeptides and activation of NOD1 in keratinocytes, while the related transporter *Slc46a3* is critical for responding to MDP, the NOD2 ligand. In a mouse model, *Slc46a2* and *Nod1* deficiency strongly suppressed psoriatic inflammation, while methotrexate, a commonly used psoriasis therapeutic, inhibited *Slc46a2*-dependent transport of DAP-muropeptides. Collectively these studies define SLC46A2 as a transporter of NOD1 activating muropeptides, with critical roles in the skin barrier, and identify this transporter as an important target for anti-inflammatory intervention.

## Introduction

Cytosolic innate immune receptors play critical roles in host defense by sensing microbial products that access the cell interior and activating potent immune and inflammatory responses (*1, 2*). These receptors include several nucleic acids sensors, such as cGAS, RIG-I, NLRP1 and AIM2, as well as others that respond to bacterial products (NOD1/2, NAIPs/NLRC4) or danger signals (NLRP1/3) (*2–7*). In some cell types, ligands for these cytosolic sensors can be imported into the cytosol (*8–12*). For instance, fragments of the bacterial cell wall, or muropeptides, enter the cytosol and drive potent inflammatory responses by activating NOD1/2 and NF-κB (*13, 14*). Muropeptides that contain the amino acid diaminopimelic acid (DAP), common to gram-negative bacteria and gram-positive bacilli, are potent NOD1 agonists, while muramyl-dipeptide (MDP), derived from nearly all bacterial peptidoglycan, activates NOD2.

While muropeptides can enter into some cell types, the underlying mechanisms of entry are unclear (*15*). Several reports have linked the solute carrier 15 (SLC15) family of peptide transporters to cytosolic muropeptide delivery (*16–21*). However, these solute carriers have not been directly demonstrated to transport muropeptides and are not specifically required for NOD signaling, but instead have recently been linked to IRF5 activation following stimulation of TLRs as well as NODs (*22, 23*). On the other hand, we recently identified the SLC46 family as candidate muropeptide transporters in *Drosophila* and mammalian cells (*24*). In particular, human or mouse *SLC46A2* or *SLC46A3*, but not *SLC46A1* (encoding the Proton Coupled Folate Transporter), strongly enhanced muropeptide-triggered NOD-dependent NF-κB reporter activity in cell lines. *SLC46A2* is particularly intriguing as it was highly effective in delivering the DAP-containing muropeptide tracheal cytotoxin (TCT) to the cytosolic innate immune receptor NOD1 in these reporter assays, yet it is expressed in only a limited set of tissues. Analysis of publicly available databases and previous publications reveals that *Slc46a2* is predominantly expressed in the skin epidermis as well as cortical epithelial cells of the thymus (*25–27*). Skin is a critical barrier defense against micro-organisms in the environment and an important immune-responsive organ (*28, 29*). NOD-mediated bacterial recognition plays a critical role in the interaction between gut microbiota and the intestinal epithelia (*30*), yet much less is known about the role of the NOD receptors in other barrier tissues, like the skin (*31*). Similar to the gut microbiome, the skin microbiome constantly interacts with the epidermis, modulating local and systemic immune responses, and is implicated in inflammatory skin diseases such as psoriasis (*32–35*). However, the role of NOD1/2 sensing in this tissue has not been evaluated.

Here we characterize a mouse deficient in *Slc46a2*, demonstrating its essential function for delivery of DAP-muropeptides and NOD1 activation in keratinocytes, and identifying a DAP-muropeptide triggered epidermal inflammatory response. This response involves Caspase-1 and Gasdermin D-dependent plasma membrane permeabilization of epidermal keratinocytes and the release of IL-1α. In the mouse, this response drives the rapid recruitment of neutrophils to muropeptide-challenged skin and is essential for the development of psoriatic inflammation. Moreover, we also show that this pathway is inhibited by the anti-folate methotrexate, indicating a novel mechanism of action for this commonly used anti-inflammatory drug (*36*). Human keratinocytes, in the context of 3D skin organoids, also similarly respond to DAP-muropeptides in an IL-1-dependent manner.

## Results

Previously, we implicated the SLC46 family of transporters in the delivery of muropeptides to cytosolic innate immune receptors in *Drosophila* and human cell lines (*24*). To determine whether SLC46 family members contribute to mammalian responses to muropeptides *in vivo*, we used a classic assay to monitor NOD1 and NOD2 activities in *Slc46a2*^-/-^ and *Slc46a3*^-/-^ mice (Fig. S1 A-E) (*37–39*). iE-DAP or MDP was injected intraperitoneally (IP) and we measured neutrophil recruitment to the peritoneum after 3 h. Like *Nod1*^-/-^ mice, *Slc46a2*^-/-^ mice did not respond to iE-DAP but responded like wild-type to MDP (Fig. 1A). By contrast, *Slc46a3*^-/-^ mice phenocopied *Nod2*^-/-^ mice, responding normally to iE-DAP but failing to respond to MDP (Fig. 1A).

**Fig. 1.**
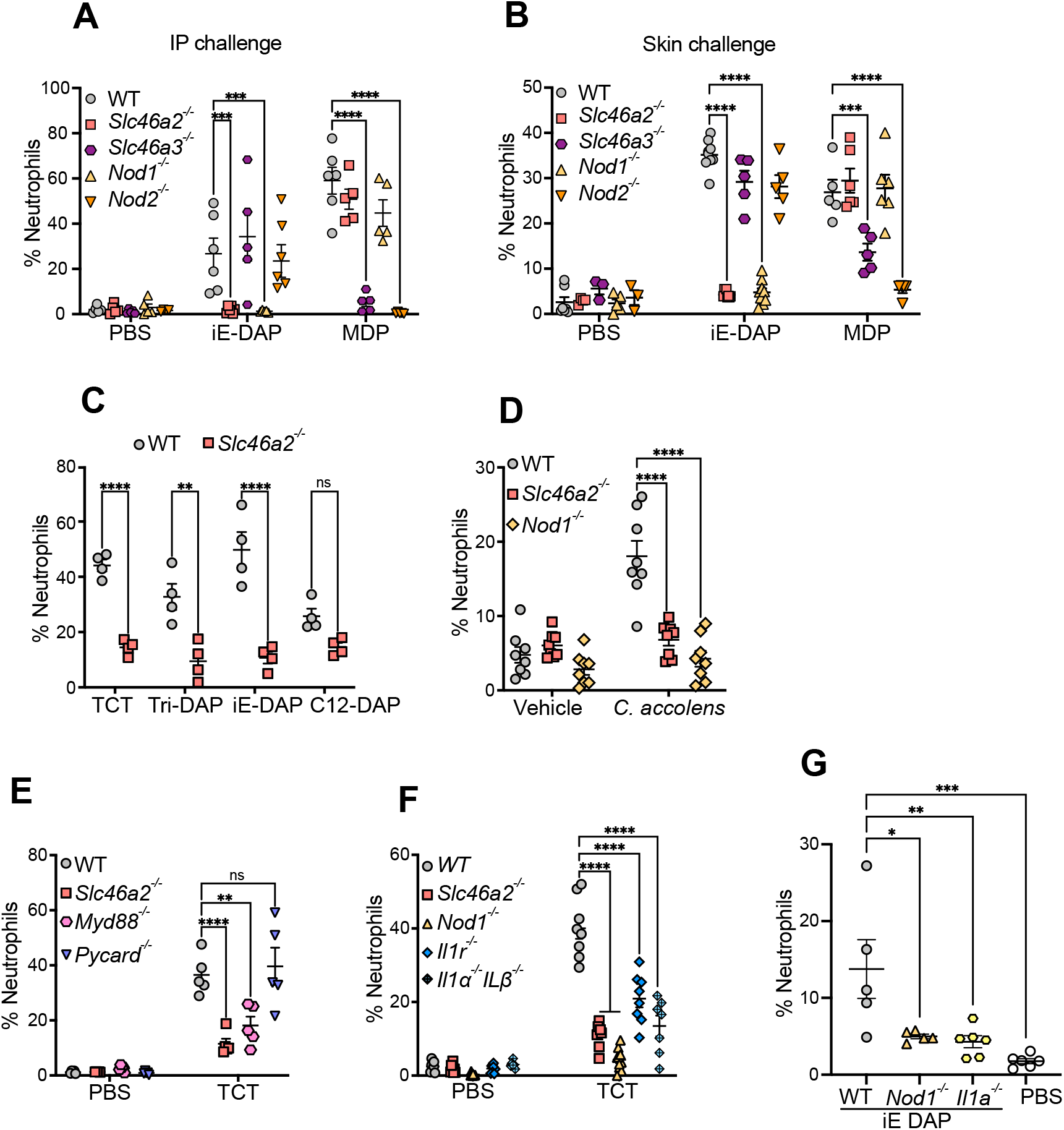
*Slc46a2* is required for neutrophil recruitment in response to NOD1 stimulation in the mouse peritoneum and skin. (**A**)and (**B**) Neutrophil recruitment after 3 h of intraperitoneal or intradermal injection of 10 μl of iE-DAP (30 μM) or MDP (10 μM). (**C**) Neutrophil recruitment after 3 h intradermal injection of different DAP-type muropeptides, TCT (8 μM), iE-DAP (30 μM), Tri-DAP (25 μM) or C12-iE-DAP (20 μM). (**D**) Neutrophil recruitment after topical association of tape stripped skin with *C. accolens*. (**E, F, & G**) Recruitment of neutrophils after 3 h of intradermal injection of 10 μl of 8 μM TCT or 30 μM iE-DAP. Genotypes indicated on all panels. Comparisons two-way ANOVA with Tukey’s multiple comparisons test to determine significance. **** P < 0.0001; *** P < 0.001; ** P < 0.01; * P < 0.05; ns, not significant.

*Slc46a2* is highly expressed in skin epidermis but not found in many other tissues (Fig. S1F and (*40, 41*)). As the activity of NOD1 agonists in skin has not been extensively characterized (*31*), the DAP-muropeptide Tracheal Cytotoxin (TCT), or LPS as a control, was injected intradermally (ID) in the pinnae and neutrophil recruitment to the ear was monitored, between 1 and 24 h (Fig. S1 G&H); leukocyte recruitment following DAP-muropeptide was rapid and robust, peaking at 3 h. Using this intradermal challenge assay, *Slc46a2*^-/-^ mice failed to respond to iE-DAP, but responded normally to MDP, phenocopying *Nod1*^-/-^ mice (Fig.1B). On the other hand, *Slc46a3*^-/-^ mice were significantly defective in responding to MDP, like *Nod2*^-/-^ animals. H&E staining of skin sections confirmed that neutrophils were recruited to the skin in response to DAP-muropeptide challenge a *Nod1*- and *Slc46a2*-dependent manner (Fig. S1I). These results, with IP or ID challenge assays, demonstrate that *Slc46a2* is selectively required for the NOD1 pathway while *Slc46a3* is selectively required for the NOD2 pathway, consistent with the idea that they each transport NOD1 or NOD2 ligands, respectively, into the cytosol.

To further probe the specificity of SLC46A2 in the response to DAP-muropeptides, we compared different NOD1 agonists (TCT, Tri-DAP, iE-DAP, and C12-iE-DAP) following ID challenge. *Slc46a2*-deficient mice displayed significantly reduced neutrophil recruitment in response to TCT, iE-DAP and Tri-DAP challenge, whereas the response to C12-iE-DAP was not significantly changed, consistent with the acyl tail enabling direct plasma membrane penetration of this molecule (Fig.1C) (*37*). Further, topical application of DAP-muropeptides on tape-stripped mouse skin also triggered robust *Slc46a2*- and *Nod1*-dependent neutrophil recruitment (Fig. S2A). Similarly, tape-stripped skin treated with live *C. accolens*, a common skin commensal with DAP-type peptidoglycan known to modulate local and systemic immunity (*32*), resulted in robust neutrophil recruitment in WT skin that was significantly decreased in either the *Slc46a2* or *Nod1* mutants (Fig. 1D). These results support the hypothesis that SLC46A2 functions in the skin epidermis to deliver DAP-muropeptides to cytosolic NOD1 in keratinocytes, triggering a rapid neutrophilic influx.

To investigate the pathways involved in DAP-muropeptide-triggered *Slc46a2/Nod1*-dependent neutrophil recruitment to the skin, *Pycard*-(encoding ASC) and *Myd88*-deficient mice were analyzed. WT and *Pycard*-deficient mice showed similar responses to ID DAP-muropeptide challenge, whereas the response was significantly reduced in *Myd88* mice, like the *Slc46a2*-deficient mice (Fig. 1E). Given that the DAP-muropeptide (TCT) used in these assays is LPS and lipopeptide free (*42*), the *MyD88* results suggest that DAP-muropeptide challenge might trigger an IL-1 response. In fact, mice lacking a functional IL-1 Receptor (*Il1r1^-/-^)* or deficient for both IL-1α and IL-1β encoding genes *(Il1a* and *Il1b)* showed significantly reduced responses to intradermal TCT challenge, similar to *Nod1^-/-^ or Slc46a2*^-/-^ mice (Fig. 1F). We further dissected the individual role of these cytokines (IL-1α and IL-1β); *Il1b*-deficient mice did not show any defect in responding to DAP-muropeptide (Fig. S2B). By contrast, *Il1a*-deficient animals failed to respond to DAP-muropeptide challenge (Fig. 1G). This phenotype was further confirmed by inhibiting lL-1α using neutralizing antibody (Fig.S2C). Keratinocytes are known for their robust IL-1α expression (*43*), and these results suggest that SLC46A2-transported DAP-muropeptide and ensuing NOD1 activation induces IL-1α release from keratinocytes, and subsequent recruitment of neutrophils in skin.

To examine IL-1α production, primary mouse keratinocytes, which express *Slc46a2* and *Nod1* (Fig.S2D), were isolated and cultured *ex vivo*, stimulated with DAP-muropeptide, and supernatants assayed for cytokine production. As predicted, WT keratinocytes released significant levels of IL-1α 24 h post challenge, whereas IL-1α released by *Slc46a2*- or *Nod1*-deficient keratinocytes was significantly reduced compared to WT, and not significantly increased compared to unstimulated cells (Fig.2A). By contrast, primary dermal fibroblasts did not respond to iE-DAP, consistent with almost no expression of *Slc46a2* and *Nod1* (Fig. S2E). Further, IP-challenge with conditioned media from DAP-muropeptide stimulated keratinocytes, but not control conditioned media, triggered significant neutrophil recruitment (Fig. S2F and 2B). However, conditioned media from IL1α-deficient keratinocytes or anti-IL-1α–depleted media from WT keratinocytes produced significantly less neutrophil attracting activity, similar to *Slc46a2*- or *Nod1*-deficient keratinocyte media (Fig. 2B and S2G). Other cytokines, including TNF, IL-6, IL-1β and IL-17 were not induced by iE-DAP challenged keratinocytes and were unchanged in the absence of *Slc46a2* or *Nod1* (Fig. S2H), while IL-23 was not detected in any condition.

**Fig. 2.**
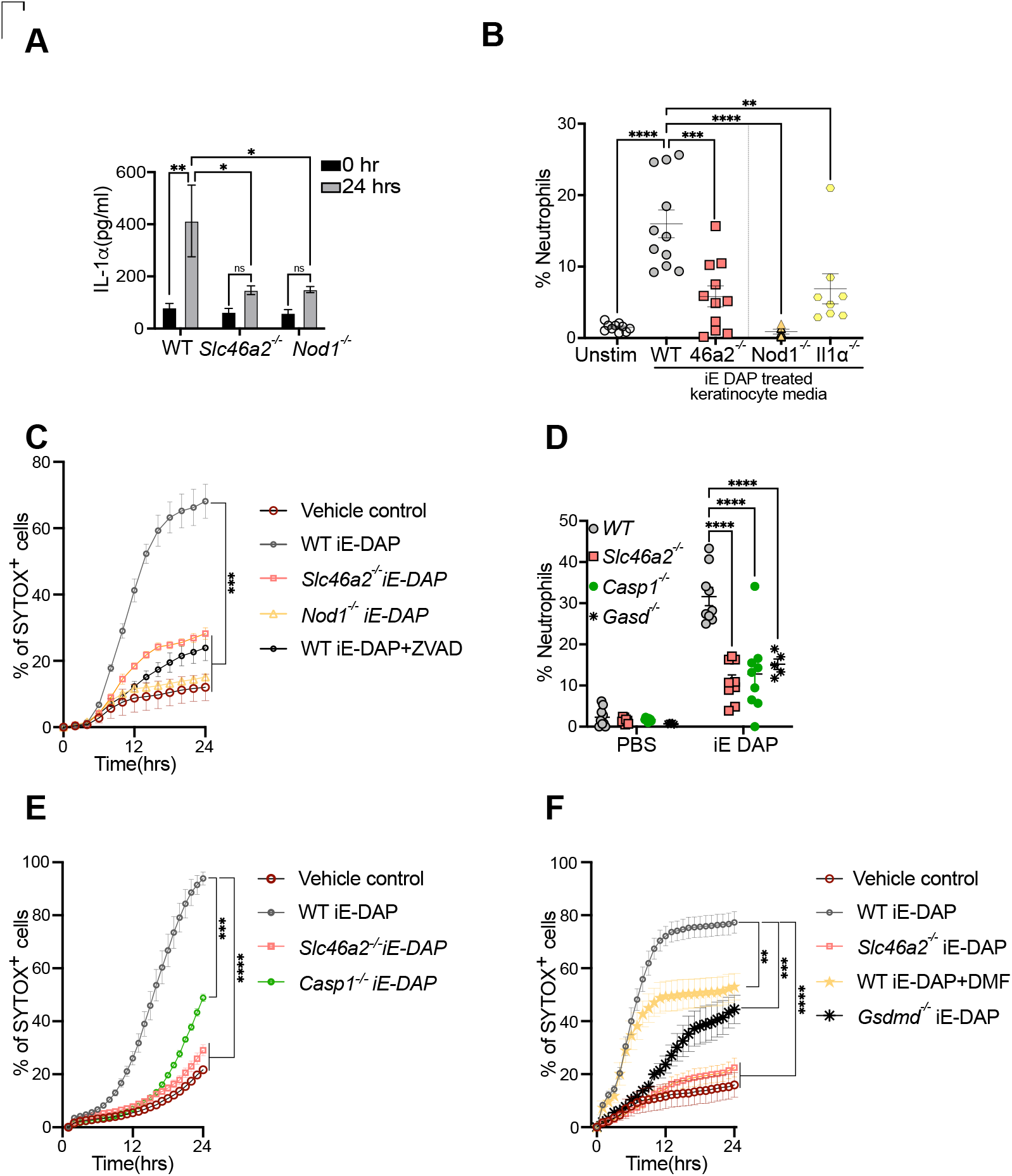
Primary mouse epidermal keratinocytes respond to DAP-muropeptides via *Slc46a2* and *Nod1*. (**A**) Keratinocytes released IL-1α following stimulation with 30 μM iE DAP for 24 h. (**B**) Neutrophils recruitment to the peritoneum after IP injection of conditioned media from WT, *Slc46a2*^-/-^, *Nod1*^-/-^ or *Il1a*^-/-^ keratinocytes stimulated with 30 μM iE-DAP for 24 h. (**C**) Sytox dye uptake by keratinocytes by live cell imaging over 24 h following 30 μM iE-DAP stimulation. (**D**) Neutrophil recruitment after 3 h intradermal injection of 10μl of 30 μM iE-DAP. (**E & F**) Sytox dye uptake by keratinocytes over 24h following 30 μM iE-DAP challenge. Genotypes, caspase inhibitor zVAD-fmk (10 μM), or GasderminD inhibitor DMF (50 μM) indicated on all panels. Panels A, B and D use two-way ANOVA and test, while C-F use one-way ANOVA, and Tukey’s multiple comparisons test to determine significance. **** P < 0.0001; *** P < 0.001; ** P < 0.01; * P < 0.05; ns, not significant. n ≥ 3 for all panels.

Alarmins, like IL-1α, are released by damaged or dying cells, triggering inflammatory responses in nearby cells (*44*). To monitor cell damage or death, we next examined DAP-muropeptide triggered keratinocyte permeabilization, quantifying membrane impermeable Sytox uptake in a live cell imaging assay(*45, 46*). WT keratinocytes were markedly permeabilized in response to DAP-muropeptide, while *Slc46a2*- and *Nod1*-deficient keratinocytes were largely protected (Fig. 2C). Interestingly, the pan-caspase inhibitor zVAD-FMK also prevented DAP-muropeptide triggered permeabilization of WT keratinocytes (Fig. 2C). To further probe the role of caspases and pyroptosis in this process, *Caspase1*- and *GasderminD*-deficient mice were challenged with ID iE-DAP injection, where they exhibited a strong defect in neutrophil recruitment, similar to *Slc46a2*^-/-^ animals (Fig 2D). Keratinocytes from these knockouts also showed significantly decreased permeabilization in response to iE-DAP (Fig. 2E & F). A similar phenotype was also observed with dimethyl fumurate (DMF), a potent inhibitor of Gasdermin pore formation (*47*). All together, these data show that DAP-muropeptide stimulation of primary keratinocytes drives cell permeabilization through a pathway requiring *Slc46a2* and *Nod1*, involving a Caspase-1/Gasdermin D pyroptosis-like process, leading to the release of IL-1α.

To directly evaluate the role of SLC46A2 in DAP-muropeptide transport, we used two complimentary chemical biological approaches. First, a modified, biologically active alkyne derivative of iE-DAP was synthesized and utilized to visualize uptake of this muropeptide into primary keratinocytes using “click-chemistry” (*48*). After 60 minutes, iE-DAP was clearly detected within both WT and *Nod1*^-/-^ cells, but not in *Slc46a2*-deficient keratinocytes, by confocal microscopy or in cell lysates (Fig. 3A and S3A, respectively). By contrast, *Slc46a2* did not affect the intracellular delivery of click-modified MDP (Fig. S3A & B), consistent with data in Figure 1 that did not implicate *Slc46a2* in the transport of this NOD2 ligand. In an orthogonal approach, fluorescently labeled lipid nanoparticles (NP) were loaded with iE-DAP and used to deliver the iE-DAP into keratinocytes, which bypassed the requirement for *Slc46a2* but not *Nod1* in inducing membrane permeability (Fig. 3B and S3C).

**Fig.3.**
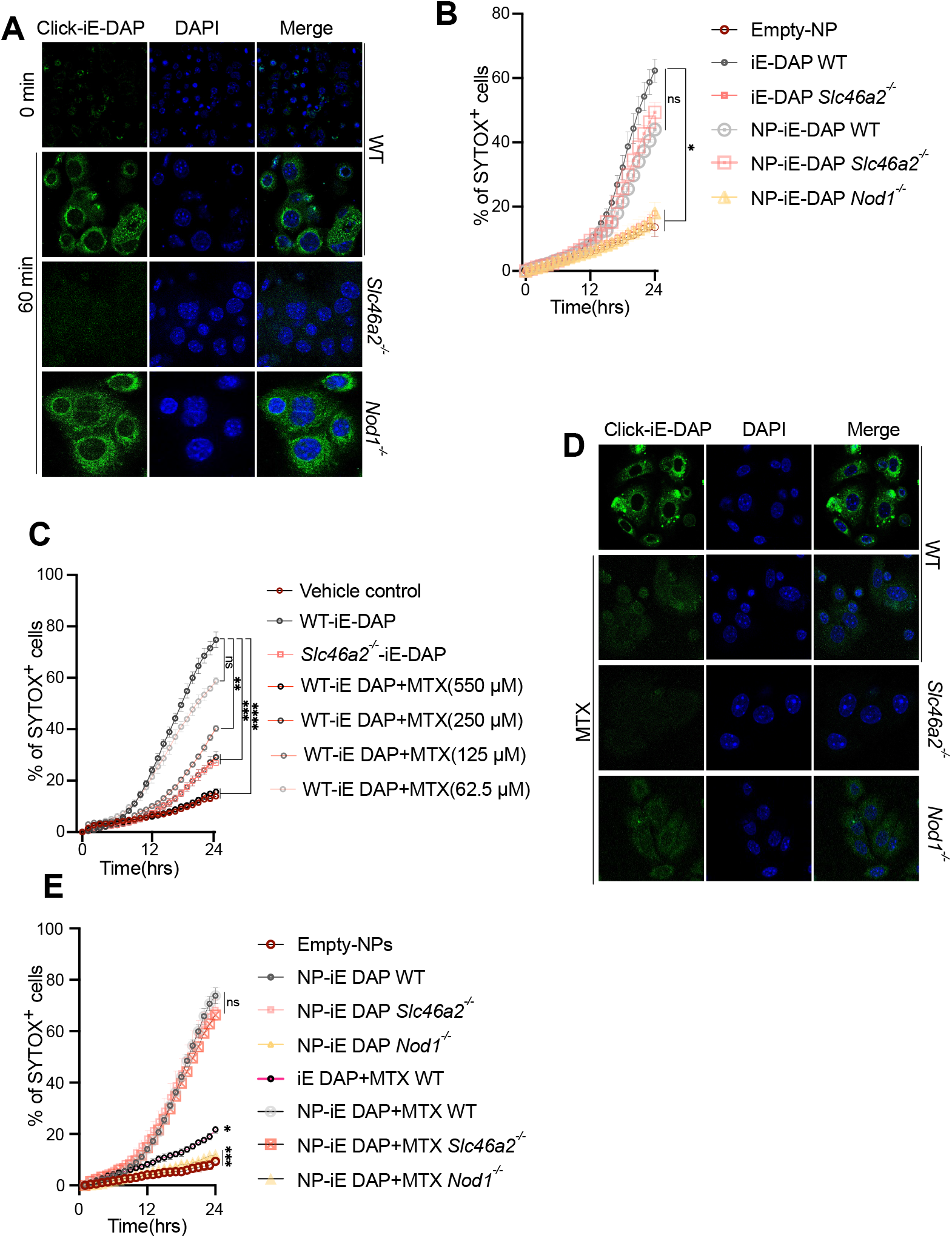
DAP-muropeptide transport requires *Slc46a2* and is blocked by methotrexate. (**A** and **D**) Confocal images of keratinocytes after 1 h challenge with 30μM “click-iE-DAP” (A) or with 30μM “click-iE DAP” and 250 μM methotrexate (MTX) (D), fixed, and then visualized with click reacted Calflour488-azide. (**B, C** and **E**) Sytox dye uptake by keratinocytes over 24h following stimulation with lipid nanoparticales (NP) loaded with iE-DAP (B), or treated increasing concentration of MTX and challenged with 30 μM iE-DAP (C), treated with 250 μM MTX and stimulated with iE-DAP-loaded NP. Genotypes indicated on all panels. Panels B, C and E uses one-way ANOVA, and Tukey’s multiple comparisons test to determine significance. **** P < 0.0001; *** P < 0.001; ** P < 0.01; * P < 0.05; ns, not significant. n ≥ 3 for all panels. Panels A and D representative images from at least three independent experiments.

SLC46A2 is a paralog of the proton-coupled folate transporter SLC46A1 (~30% identity), which suggests folates and anti-folates could be a common cargo for all SLC46 family proteins (*49, 50*). The anti-folate methotrexate (MTX) is a potent anti-inflammatory drug commonly used to treat psoriasis and rheumatoid arthritis, with unclear mechanisms of action (*36, 51*). This led us to hypothesize that MTX competes with DAP-muropeptides for docking to/transport by SLC46A2, which was tested by adding increasing concentrations of MTX in the keratinocyte-based iE-DAP assays. In a dose dependent manner, MTX inhibited *Slc46a2*-dependent DAP-muropeptide triggered keratinocyte permeabilization and IL-1α release, similar to the *Slc46a2*-deficient keratinocytes (Fig. 3C and S3D). Further, MTX blocked the cytosolic accumulation of “click”-iE-DAP, similar to competition with unlabeled iE-DAP (Fig. 3D and S3E) but failed to interfere with NP-mediated iE-DAP delivery (Fig. 3F and S3F).

The above results show that SLC46A2 is inhibited by MTX and is required for a skin inflammatory response to *C. accolens.* Interestingly, MTX is used as a first line treatment for the inflammatory skin disease psoriasis, while *Corynebacterium spp.* are linked to psoriasis and known to exacerbate psoriasis-like phenotypes in a mouse model (*33–36*). Therefore, we used the imiquimod (IMQ) model to probe the role of *Slc46a2* and *Nod1* in psoriatic-like inflammation. With a 7-day course of topical IMQ application, WT mice displayed the expected psoriatic-like inflammation, assayed by enhanced ear thickness and H&E histology, while *Slc46a2*- and *Nod1*-deficient mice were markedly resistant (Fig. 4A, B and S4A). Application of IMQ for only 3 days followed by topical application *C. accolens* similarly drove psoriasis-like inflammation in WT mice while both *Slc46a2* and *Nod1* deficient animals presented dramatically reduced inflammation (Fig.4C and S4B, C). Further, topical application of MTX reduced the psoriasis-like inflammation in WT skin, to levels similar to the mutant strains, while MTX had no observable effect on the residual inflammation in *Slc46a2*^-/-^ and *Nod1*^-/-^ skin (Fig. 4D and S4D, E).

**Fig. 4.**
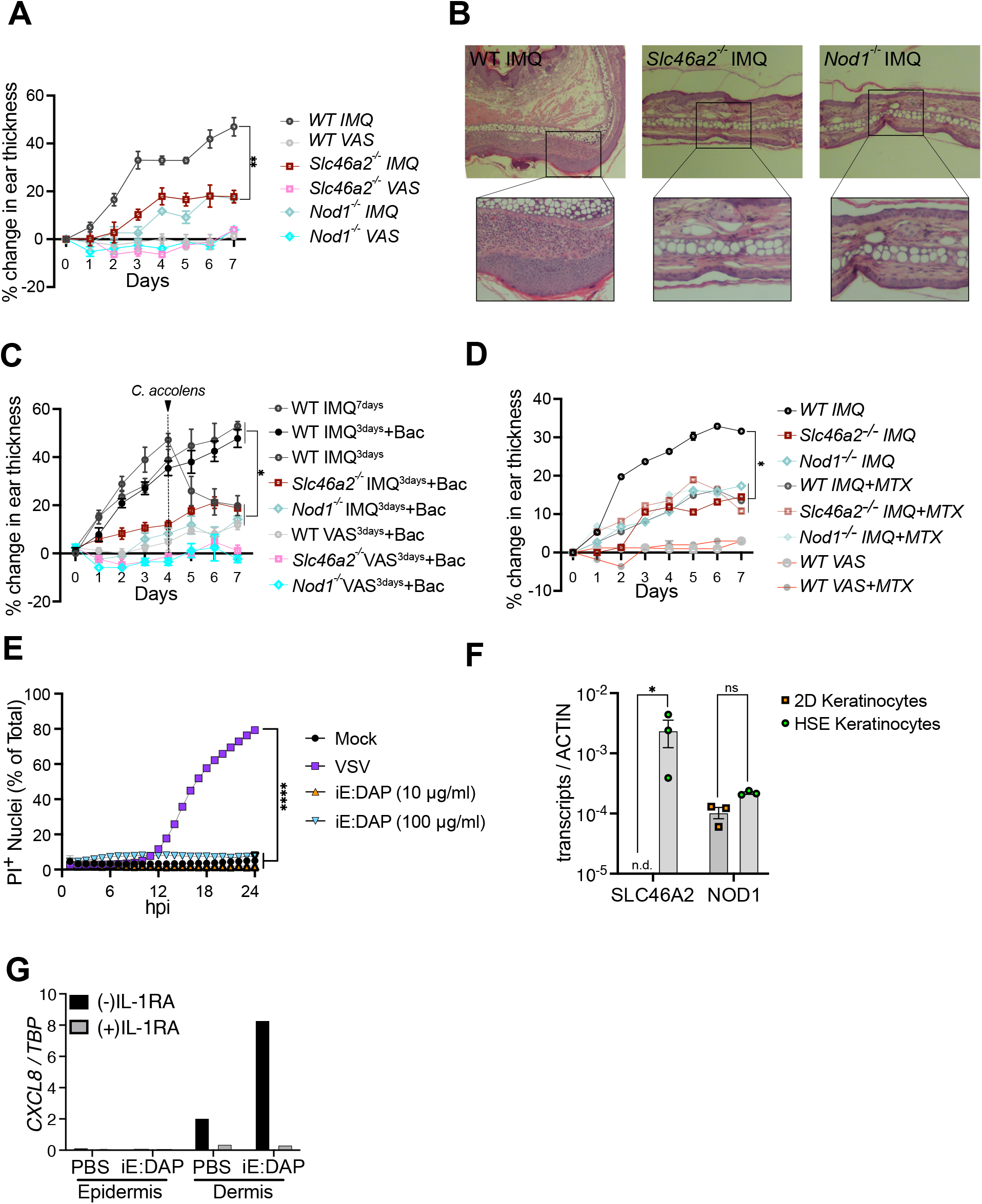
*Slc46a2*^-/-^ and *Nod1*^-/-^ mice are resistant to IMQ-induced psoriatic inflammation. (**A, C** and **D**) 5% Imiquimod (IMQ) was topically applied to pinnae daily to induce psoriasis and ear inflammation was quantified daily. Contralateral pinnae were treated with Vaseline (VAS) as vehicle control. For (A), mice were treated with IMQ daily for 7 days and mean pinnae thickness is plotted. (**B**) H&E stained histology images from Imiquimod treated pinnae on day 7. Genotypes are indicated on all panels. (**C**) IMQ was applied for only 3 days, and then pinnae were treated daily for 3 days with topical application of live *C. accolens* (10^7^ CFU), except for controls with either a full 7 days or just 3 days of IMQ treatment. (**D**) is similar to (A) except IMQ was applied daily along with xx volument of 250 μM MTX. (**E**) Propidium Iodide (PI) uptake assay using primary human foreskin keratinocytes challenged with indicated iE-DAP doses or VSV virus infection (MOI 10) as a positive control. iE-DAP treatment did not induce cell permeabilization in human keratinocytes. (**F**) Expression analysis of *SLC46A2* and *NOD1* in keratinocytes grown in 2D culture and 3D organoids (HSE). (**G**) Induction of *CXCL8* in skin organoid epidermal and dermal layers after PBS or iE-DAP challenge in the presence and absence of IL-1 receptor inhibitor (IL-1RA). High expression of *CXCL8* was observed in dermis following iE-DAP challenge, and this was blocked by IL-1RA. Panels A, C, D and E use one-way ANOVA and Panel F uses two-way ANOVA test. Tukey’s multiple comparisons test was used to determine significance. Panel B and G are representative of three independent experimental results. **** P < 0.0001; *** P < 0.001; ** P < 0.01; * P < 0.05; ns, not significant. n ≥ 3 for all panels.

To determine if iE-DAP-triggered inflammatory responses occur in human skin, we first analyzed primary foreskin-derived human keratinocytes. For this experiment, keratinocytes were infected with vesicular stomatitis virus (VSV) as a positive control, as it is known to induce cell permeabilization via a GSDME-dependent pyroptotic pathway (Fig. 4E) (*46*). By contrast, iE-DAP treatment failed to induce membrane permeabilization in these cells (Fig. 4E). Interestingly, primary human keratinocytes, in standard tissue culture conditions, did not express *SLC46A2* (Fig 4F). When these same keratinocytes are cultured at an air-liquid interface on a bed of collagen-embedded dermal fibroblasts they differentiate into a multilayered squamous epithelium (known as a Human Skin Equivalents (HSE)), with many similarities to human skin (*52*). In the context of the HSE, differentiated human keratinocytes robustly expressed *SLC46A2* and these organoids responded to iE-DAP. Upon treatment of the HSE epidermis, but not the dermis, with 30 μM iE-DAP for 24 h, we observed the induction of *CXCL8* in dermal fibroblasts (Fig. 4G and S4F). Moreover, this response was prevented by concomitant treatment with recombinant IL-1R antagonist (IL-1Ra), strongly implicating IL-1 release by keratinocytes in driving *CXCL8* induction in HSE dermal fibroblasts. Likewise, in the mouse system, supernatants from iE-DAP-stimulated primary keratinocytes triggered, via *Slc46a2*- and *Nod1*-dependent signaling, CXCL1 (KC) production in dermal fibroblasts (S4G).

## Discussion

While it has been clear for many years that some cell types import muropeptides and activate the cytosolic innate immune receptors NOD1 or NOD2 (*53*), the underlying cellular and molecular mechanisms that mediate import of these molecules has remained unclear. Initially, the SLC15 family of oligopeptide transporters was implicated in this process (*16–20*), but they have never been demonstrated to bind or transport muropeptides while more recent studies instead have shown SLC15A4 functions as a scaffold involved in IRF5 activation downstream of TLR as well as NOD activation (*20, 22*). On the other hand, our initial work in *Drosophila* implicated SLC46s in this process (*24*), and here we show that SLC46A2 has the properties of a transporter selective for delivering DAP-type muropeptides for NOD1 activation in murine tissues. Using *in vivo* approaches in two different tissues, as well as *ex vivo* studies with primary mouse keratinocytes, *Slc46a2* was uniquely required for the response to multiple NOD1 activating DAP-muropeptides, while *Slc46a3* was required for the response to the NOD2 activating muropeptide MDP. Moreover, two orthogonal approaches were used to demonstrate that intracellular delivery of DAP-muropeptide requires *Slc46a2.* Intracellular iE-DAP, directly detected via click chemistry dye labeling, required *Slc46a2* but not *Nod1.* On the other hand, when packaged in a lipo-nanoparticle for direct intracellular delivery, iE-DAP activation of NOD1 no longer required *Slc46a2.* Together, these immunological and cell biological data strongly argue for the direct transport of DAP-muropeptides by SLC46A2, and implicate SLC46A3 in the delivery of MDP, and not vice versa. Interestingly, primary human keratinocytes, in standard tissue culture conditions, do not express *SLC46A2* and do not respond to iE-DAP, while after differentiating into a squamous epithelium in the HSE organoids, *SLC46A2* is expressed and the epidermis responds to DAP-muropeptide.

In both humans and mice, *Slc46a2* is expressed in only a limited set of tissues, including the epidermis (*40, 41*). This expression pattern led us to explore, in more detail, the role of SLC46A2/NOD1 dependent responses in skin. We found a robust neutrophilic response to intradermal DAP-muropeptide challenge that required *Slc46a2* and *Nod1.* Similarly, we also found that murine skin, once the outer waxy stratum corneum was removed, responded to topical application of either iE-DAP or the DAP-producing skin commensal *Corynebacterium accolens* through *Slc46a2* and *Nod1.* Likewise, primary mouse keratinocytes *ex vivo* responded to DAP-muropeptides with cell permeabilization and the release of active IL-1α. Similar to the *in vivo* neutrophilic response, the cell permeabilization pathway in keratinocytes involved *Casp1* and *Gsdmd* in addition to *Slc46a2* and *Nod1*. Together these results demonstrate that skin keratinocytes respond to DAP-muropeptides via intracellular delivery by SL46A2 and activation of NOD1, which in turn drives IL-1α release via pathway that involves Caspase-1 and Gasdermin D cell permeabilization, and perhaps cell death. The rapidity of the response *in vivo* suggests this is not a transcription-dependent response, although the response is slower in isolated keratinocytes.

Given the established link between *C. accolens* and psoriasis, in humans and mouse models (*33–35*), the role of this response in psoriatic inflammation was also examined in a mice model. Animals deficient for either *Slc46a2* or *Nod1* were strikingly resistant to psoriatic inflammation and unresponsive to *C. accolens* triggered pathology. Given the previous work linking *C. accolens* to the activation of IL-17 producing γδT cells in the skin (*32*), it will be interesting to learn if SLC46A2/NOD1 pathway in the skin also drives this IL-17 producing γδ response, in addition to the strong neutrophilic infiltration.

Immune-mediated inflammatory diseases, like psoriasis, are often treated with the first line drug methotrexate. MTX is a complicated drug, originally developed as an antiproliferative due to its interference with folate-dependent enzymes required for nucleotide biosynthesis and DNA replication, that is still used as a cancer therapy. Subsequently, MTX was found to be effective in psoriasis, rheumatoid arthritis and other immune mediate inflammatory diseases, but at much lower doses that do not impact cell proliferation(*36, 54*). The mechanisms of action of MTX as an anti-inflammatory are controversial. The leading model argues that MTX functions as a traditional anti-folate interfering with the enzyme AICAR transformylase, leading to increased intracellular AICAR and eventually increased extracellular Adenosine, a known anti-inflammatory molecule that acts via adenosine receptors, such as P1-A_2A_ (*55*). In addition, MTX has been argued to interfere with NF-κB activation through unknown mechanisms. Here we present a new mechanism of action for anti-inflammatory activity of MTX that does not involve interfering with folate-dependent enzymes. Instead, the data here shows that MTX competes with DAP-muropeptide for delivery into the cytosol, phenocopying *Slc46a2*-deficiency in three separate *ex vivo* keratinocytes assays as well as with a *in vivo* model of psoriatic inflammation. *In vivo*, MTX application in the *Slc46a2 and Nod1* mutants had no effect on the residual inflammation still observed in the psoriasis model, while it strongly suppressed psoriatic-like inflammation in WT mice, to levels very similar to that observed in the mutants. Together, these *in vitro* and *in vivo* findings argue that MTX competes with DAP-muropeptides as cargo for the transporter SLC46A2, and thereby directly interferes with a central inflammatory pathway, the NOD1 pathway, which operates in the skin to drive neutrophil recruitment and psoriatic-like inflammation. In the future, biophysical approaches will be necessary to probe the cargo binding activity and specificity of SLC46A2, as well as the related transporters SLC46A1 (the established proton-coupled folate/anti-folate transporter) and SLC46A3; all three members of the SLC46 family share approximately 30% amino acid identity. Future work will also focus on precisely mapping the anatomical location(s) of these responses within the epidermis and probe the connection between SLC46A2/NOD1-mediated response in keratinocytes to activation of IL-17 producing γδ T-cells in the skin, which are known to contribute to psoriasis in mouse models and humans and are activated by *C. accolens* (*32, 56*).

Through a combination of immunology, chemical biology and biomedical engineering approaches, we demonstrate that SLC46A2 functions to deliver DAP-muropeptides to the cytosol of keratinocytes, driving a robust IL-1α dependent inflammatory response and neutrophil recruitment. This response is critical for psoriatic-like inflammation in a mouse model, while human skin organoids respond similarly although the connection to psoriasis awaits future analysis. Additionally, the common anti-inflammatory drug MTX phenocopies *Slc46a2* deficiency *in vivo* and in primary mouse keratinocytes, arguing that SLC46A2 is a direct anti-inflammatory target of this drug. Moreover, these findings identify disease mechanisms and therapeutic targets that could be applicable to a broad number of inflammatory conditions and highlight a novel role for the SLC46 family in host-microbiome interactions.

## Acknowledgments

The authors wish to thank Douglas Golenbock, Kate Fitzgerald, Egil Lien and Gabriel Nunez for sharing mouse strains, and Egil Lien for comments on the manuscript, Gail German and Zhaozhao Jiang for technical assistance.

## Funding

National Institutes of Health grant ROAI060025 (NS)

Kenneth Rainin Foundation (NS)

National Institutes of Health grant 5T32GM133395 (KW)

National Institutes of health grant R01GM138599 (CLG)

Pew Biomedical scholars Program (CLG)

## Author contributions

Conceptualization: DP, MO, AUP, CLG, NS

Methodology: RB, CL, SM, AN, MS, GIK, KW, AB, KO, AM, AN, MA, WEG, TDK, CLG, PUA, NS

Investigation: RB, AN, MS, KO, PR

Funding acquisition: NS, CLG

Supervision: NS

Writing – original draft: RB, NS

Writing – review & editing: NS

## Competing interests

A provisional patent on targeting SLC46s to inhibit inflammation in psoriasis and other auto-inflammatory diseases as been filed by some of the authors.

## Data and materials availability

All data are available in the main text or the supplementary materials. Mouse strains and other reagents will be shared with academic researchers with a simple MTA.

## Supplementary Materials

## Materials and Methods

## Bibliography

1. S. A. Ragland, J. C. Kagan, Cytosolic detection of phagosomal bacteria-Mechanisms underlying PAMP exodus from the phagosome into the cytosol. Mol Microbiol 116, 1420–1432 (2021).

2. V. A. Rathinam, K. A. Fitzgerald, Cytosolic surveillance and antiviral immunity. Curr Opin Virol 1, 455–462 (2011).

3. H. Okude, D. Ori, T. Kawai, Signaling Through Nucleic Acid Sensors and Their Roles in Inflammatory Diseases. Front Immunol 11, 625833 (2020).

4. K. V. Swanson, M. Deng, J. P. Ting, The NLRP3 inflammasome: molecular activation and regulation to therapeutics. Nat Rev Immunol 19, 477–489 (2019).

5. E. L. Johnston, B. Heras, T. A. Kufer, M. Kaparakis-Liaskos, Detection of Bacterial Membrane Vesicles by NOD-Like Receptors. Int J Mol Sci 22, (2021).

6. B. Sundaram, T. D. Kanneganti, Advances in Understanding Activation and Function of the NLRC4 Inflammasome. Int J Mol Sci 22, (2021).

7. S. Bauernfried, M. J. Scherr, A. Pichlmair, K. E. Duderstadt, V. Hornung, Human NLRP1 is a sensor for double-stranded RNA. Science 371, (2021).

8. M. Kudo, K. Kobayashi-Nakamura, N. Kitajima, K. Tsuji-Naito, Alternate expression of PEPT1 and PEPT2 in epidermal differentiation is required for NOD2 immune responses by bacteria-derived muramyl dipeptide. Biochem Biophys Res Commun 522, 151–156 (2020).

9. R. D. Luteijn et al., SLC19A1 transports immunoreactive cyclic dinucleotides. Nature 573, 434–438 (2019).

10. C. Ritchie, A. F. Cordova, G. T. Hess, M. C. Bassik, L. Li, SLC19A1 Is an Importer of the Immunotransmitter cGAMP. Mol Cell 75, 372–381 e375 (2019).

11. W. Strober, T. Watanabe, NOD2, an intracellular innate immune sensor involved in host defense and Crohn’s disease. Mucosal Immunol 4, 484–495 (2011).

12. Z. Al Nabhani, G. Dietrich, J. P. Hugot, F. Barreau, Nod2: The intestinal gate keeper. PLoS Pathog 13, e1006177 (2017).

13. K. H. Schleifer, O. Kandler, Peptidoglycan types of bacterial cell walls and their taxonomic implications. Bacteriol Rev 36, 407–477 (1972).

14. T. Mukherjee et al., NOD1 and NOD2 in inflammation, immunity and disease. Arch Biochem Biophys 670, 69–81 (2019).

15. O. Irazoki, S. B. Hernandez, F. Cava, Peptidoglycan Muropeptides: Release, Perception, and Functions as Signaling Molecules. Front Microbiol 10, 500 (2019).

16. M. G. Ismair et al., hPepT1 selectively transports muramyl dipeptide but not Nod1-activating muramyl peptides. Can J Physiol Pharmacol 84, 1313–1319 (2006).

17. G. M. Charriere et al., Identification of Drosophila Yin and PEPT2 as evolutionarily conserved phagosome-associated muramyl dipeptide transporters. J Biol Chem 285, 20147–20154 (2010).

18. J. Lee et al., pH-dependent internalization of muramyl peptides from early endosomes enables Nod1 and Nod2 signaling. J Biol Chem 284, 23818–23829 (2009).

19. N. Nakamura et al., Endosomes are specialized platforms for bacterial sensing and NOD2 signalling. Nature 509, 240–244 (2014).

20. S. Sasawatari et al., The solute carrier family 15A4 regulates TLR9 and NOD1 functions in the innate immune system and promotes colitis in mice. Gastroenterology 140, 1513–1525 (2011).

21. Y. Hu, F. Song, H. Jiang, G. Nunez, D. E. Smith, SLC15A2 and SLC15A4 Mediate the Transport of Bacterially Derived Di/Tripeptides To Enhance the Nucleotide-Binding Oligomerization Domain-Dependent Immune Response in Mouse Bone Marrow-Derived Macrophages. J Immunol 201, 652–662 (2018).

22. L. X. Heinz et al., TASL is the SLC15A4-associated adaptor for IRF5 activation by TLR7-9. Nature 581, 316–322 (2020).

23. I. Rimann et al., The solute carrier SLC15A4 is required for optimal trafficking of nucleic acid-sensing TLRs and ligands to endolysosomes. Proc Natl Acad Sci U S A 119, e2200544119 (2022).

24. D. Paik et al., SLC46 Family Transporters Facilitate Cytosolic Innate Immune Recognition of Monomeric Peptidoglycans. J Immunol 199, 263–270 (2017).

25. C. Chen et al., Characterization of the mouse gene, human promoter and human cDNA of TSCOT reveals strong interspecies homology. Biochim Biophys Acta 1493, 159–169 (2000).

26. K. Y. Kim et al., Expression Analyses Revealed Thymic Stromal Co-Transporter/Slc46A2 Is in Stem Cell Populations and Is a Putative Tumor Suppressor. Mol Cells 38, 548–561 (2015).

27. M. Uhlen et al., Proteomics. Tissue-based map of the human proteome. Science 347, 1260419 (2015).

28. P. M. Elias, Skin barrier function. Curr Allergy Asthma Rep 8, 299–305 (2008).

29. J. A. Bouwstra, M. Ponec, The skin barrier in healthy and diseased state. Biochim Biophys Acta 1758, 2080–2095 (2006).

30. K. S. Kobayashi et al., Nod2-dependent regulation of innate and adaptive immunity in the intestinal tract. Science 307, 731–734 (2005).

31. J. Harder, G. Nunez, Functional expression of the intracellular pattern recognition receptor NOD1 in human keratinocytes. J Invest Dermatol 129, 1299–1302 (2009).

32. V. K. Ridaura et al., Contextual control of skin immunity and inflammation by Corynebacterium. J Exp Med 215, 785–799 (2018).

33. A. Balato et al., Human Microbiome: Composition and Role in Inflammatory Skin Diseases. Arch Immunol Ther Exp (Warsz) 67, 1–18 (2019).

34. Z. Stehlikova et al., Crucial Role of Microbiota in Experimental Psoriasis Revealed by a Gnotobiotic Mouse Model. Front Microbiol 10, 236 (2019).

35. P. Zanvit et al., Antibiotics in neonatal life increase murine susceptibility to experimental psoriasis. Nat Commun 6, 8424 (2015).

36. A. K. J. a. M. E. Weinblatt, in Rheumatology, M. H. A. S. E. G. J. S. M. W. M. Weisman, Ed. (Elsevier, Philadelphia PA USA, 2018), vol. 1, chap. 66.

37. J. Masumoto et al., Nod1 acts as an intracellular receptor to stimulate chemokine production and neutrophil recruitment in vivo. J Exp Med 203, 203–213 (2006).

38. H. L. Rosenzweig et al., Activation of NOD2 in vivo induces IL-1beta production in the eye via caspase-1 but results in ocular inflammation independently of IL-1 signaling. J Leukoc Biol 84, 529–536 (2008).

39. J. G. Magalhaes et al., Murine Nod1 but not its human orthologue mediates innate immune detection of tracheal cytotoxin. EMBO Rep 6, 1201–1207 (2005).

40. C. Wu, X. Jin, G. Tsueng, C. Afrasiabi, A. I. Su, BioGPS: building your own mash-up of gene annotations and expression profiles. Nucleic Acids Res 44, D313–316 (2016).

41. M. Karlsson et al., A single-cell type transcriptomics map of human tissues. Sci Adv 7, (2021).

42. T. Kaneko et al., Monomeric and polymeric gram-negative peptidoglycan but not purified LPS stimulate the Drosophila IMD pathway. Immunity 20, 637–649 (2004).

43. M. H. Orzalli et al., An Antiviral Branch of the IL-1 Signaling Pathway Restricts Immune-Evasive Virus Replication. Mol Cell 71, 825–840 e826 (2018).

44. N. C. Di Paolo, D. M. Shayakhmetov, Interleukin 1alpha and the inflammatory process. Nat Immunol 17, 906–913 (2016).

45. P. Orning et al., Pathogen blockade of TAK1 triggers caspase-8-dependent cleavage of gasdermin D and cell death. Science 362, 1064–1069 (2018).

46. M. H. Orzalli et al., Virus-mediated inactivation of anti-apoptotic Bcl-2 family members promotes Gasdermin-E-dependent pyroptosis in barrier epithelial cells. Immunity 54, 1447–1462 e1445 (2021).

47. F. Humphries et al., Succination inactivates gasdermin D and blocks pyroptosis. Science 369, 1633–1637 (2020).

48. P. Shieh et al., CalFluors: A Universal Motif for Fluorogenic Azide Probes across the Visible Spectrum. J Am Chem Soc 137, 7145–7151 (2015).

49. J. L. Parker et al., Structural basis of antifolate recognition and transport by PCFT. Nature 595, 130–134 (2021).

50. R. Zhao, S. Aluri, I. D. Goldman, The proton-coupled folate transporter (PCFT-SLC46A1) and the syndrome of systemic and cerebral folate deficiency of infancy: Hereditary folate malabsorption. Mol Aspects Med 53, 57–72 (2017).

51. A. M. Alqarni, M. P. Zeidler, How does methotrexate work? Biochem Soc Trans 48, 559–567 (2020).

52. M. W. Carlson, A. Alt-Holland, C. Egles, J. A. Garlick, Three-dimensional tissue models of normal and diseased skin. Curr Protoc Cell Biol Chapter 19, Unit 19 19 (2008).

53. P. A. D. Bastos, R. Wheeler, I. G. Boneca, Uptake, recognition and responses to peptidoglycan in the mammalian host. FEMS Microbiol Rev 45, (2021).

54. E. Schrezenmeier, T. Dorner, Mechanisms of action of hydroxychloroquine and chloroquine: implications for rheumatology. Nat Rev Rheumatol 16, 155–166 (2020).

55. B. N. Cronstein, T. M. Aune, Methotrexate and its mechanisms of action in inflammatory arthritis. Nat Rev Rheumatol 16, 145–154 (2020).

56. P. Rider et al., IL-1alpha and IL-1beta recruit different myeloid cells and promote different stages of sterile inflammation. J Immunol 187, 4835–4843 (2011).

