## Supplementary Methods and Figures for "Methotrexate inhibition of muropeptide transporter SLC46A2 controls psoriatic skin inflammation"

##### Ethics

All animal studies were performed in compliance with the federal regulations set forth in the Animal Welfare Act (AWA), the recommendations in the Guide for the Care and Use of Laboratory Animals of the National Institutes of Health, and the guidelines of the UMass Medical School Institutional Animal Use and Care Committee. All protocols used in this study were approved by the Institutional Animal Care and Use Committee at the UMass Medical School (protocol 2056).

##### Mice

ES cells harboring *Slc46a2* locus targeted with Zen-Ub1 cassette were purchased from Mutant Mouse Resource and Research Center (MMRRC) at University of California at Davis, an NIH-funded strain repository, and were donated to the MMRRC by The KOMP Repository, University of California, Davis; originating from David Valenzuela, George Yancopoulos, Regeneron Pharmaceuticals, Inc (RRID:MMRRC_062453-UCD) (*1*). ES cells were used to generate chimeric founder mice by standard microinjection into albino C57BL/6J blastocysts, by the UMMS transgenic mouse core. Chimeras were mated to albino C57Bl/6J mice to identify germline transmission. Heterozygous animals were identified using PCR (see Key Resources Table for oligos). Wildtype (*Slc46a2*$^{+/+}$) and mutant (*Slc46a2*$^{-/-}$) lines were established from these heterozygous and used in all experiments. *Slc46a3*$^{-/-}$ mice (*Slc46a3*$^{tm1.1(KOMP)Vlcg}$) were generated by KOMP-Regeneron and acquired from Jackson Labs.

For *in vivo* analysis of NOD1/2 responses in mouse skin, mixed male and female mice aged 4-6 weeks were used, while for IP NOD1/2 challenge assays male mice aged 6-12 weeks were used.

For genotyping, mice were anesthetized, and an ear-punch or tail sample collected in a sterile blade in a PCR tube. 75 µl of DNA extraction buffer-75ul (25mM NaOH and 0.2 mM EDTA) was added to sample, and it was heated at 95°C for 1hr in a thermocycler. Samples was then treated 75 µl of neutralizing buffer (40mM Tris Ph-5.5) and stored at analyzed by PCR. Primer sequences are reported in the KRT.

*Nod1*$^{-/-}$, *Nod2*$^{-/-}$, *IL1a*$^{-/-}$, *IL1b*$^{-/-}$, *IL1R*$^{-/-}$, *Pycard*$^{-/-}$, *and MyD88*$^{-/-}$, *Casp1*$^{-/-}$, and *Gasmd*$^{-/-}$ mice were reported previously (*2-9*) . Note that these Caspase-1 knockouts are wild-type for Caspase-11.

##### Keratinocyte isolation, culture and analysis

Adult (6-8 weeks) mouse keratinocytes were isolated from male or female animals using protocols described previously (*10*). In brief, tail skin was incubated in 1mg/ml Dispase II (Roche) overnight at 4°C. The epidermis was physically removed from the dermis with tweezers following Dispase II treatment, subsequently digested with TrypLE (Thermo) for 20min at room temperature, and keratinocytes were detached by vigorously shaking in culture medium and filtered with a 70 µm strainer (Fisher scientific). Isolated cells were seeded at a density of $10^5$cells/cm$^2$ cultured with EpiLife (Gibco) in 12-well plates precoated with coating matrix (Gibco) and used between 3 and 5 days after isolation.

For preparation of conditioned media, keratinocytes were first stimulated with 8 μM TCT, 30 μM iE-DAP or 20ng/ml LPS for 1 hour (or left unstimulated). Then cells were washed 3 times with culture medium to remove agonists and cultured further in complete media for 24 hours, when the media was collected and centrifuged. 200μl of conditioned media was intraperitoneally injected into naïve mice and leukocyte recruitment assayed, as below.

For cytokine ELISAs, primary keratinocytes were challenged, or not, with 8 μM TCT or 30 μM iE-DAP for 24 h and culture medium was harvested. Media was centrifuged at 500g for 5mins and used for analyzing cytokine levels by ELISA (R&D systems) using manufacturer's protocol. In brief, 96-well ELISA plate were coated with capture antibody overnight at room temperature. The next day, the plate was washed three times with wash buffer and incubated with media (100μl) for 2h followed by three washes and then probed with biotin-labelled detection antibody for 2h. After three washes plated were incubated with HRP-streptavidin, developed with TMB (3,3', 5,5"-tetramethylbenzidine) substrate for 20 min at room temperature, and then stopped with 1N sulfuric acid. Plate was analyzed at 450 nm with wavelength correction at 540 nm.

For cell permeabilization assays, $1X10^5$ keratinocytes /well were seeded in a 96-well plate (Denville scientific inc.). After reaching 80% confluency 30 μM iE-DAP was added with 1nM Sytox red dye (Thermo) and 0.2nM Hoechst stain (Thermo). The plate was then cultured in the Cytation5 (BioTek) live-imaging microscope with quantitive imaging of 1 field of view in each well every 60 min for 24 hours. For analysis, total number of intact cell nuclei were quantified via Hoechst positive nuclei and total number of permeabilized cells were quantified by Sytox positive nuclei. Percentage of $Sytox^+$ cells was then calculated.

**Intradermal pinnae injections**

10μl of 30 μM iE DAP, 8μM TCT, 25μM Tri-DAP, 20μM C12-iE DAP or PBS was injected in ventral side of right and left pinnae, respectively. After indicated time, both ears were individually harvested, stored in ice cold PBS on ice until further processing for flow cytometry staining.

**Intraperitoneal injections**

500μl of iE DAP (30μM) or MDP (20μM) or PBS were intraperitoneal injected for 3h and peritoneal cells were harvested from euthanized mice by injecting 8ml of ice-cold PBS (*11*). Harvested cell suspensions were centrifuged at 300g for 5 min at 4°C, washed in PBS, and resuspended in FACS buffer.

**Pinnae processing**

Pinnae were cut into small pieces and incubated for 30 min at 37°C in a solution of 1mg/ml Dispase-II (Roche) in DMEM media containing 10% FBS. Ice-cold FACS buffer (2% FBS in PBS) was added to skin samples to stop the reaction. Then samples were placed on 70μm cell strainers and ground using cell strainer pestle to create a single cell suspension. Cells were spun at 180g for 5min at 4°C and resuspended in FACS buffer.

**Flow cytometry**

For skin,1X10$^6$ cells were incubated in Fc receptor block (anti CD16) for 10 min and stained with F4/80-PE/Cy7 (1:200), CD45-alexaflour700 (1:200) and Gr1-PE (1:200) for 30 min in ice in the dark. Samples were washed thrice in FACS buffer and analyzed with a Cytek Aurora cytometer. For peritoneal cells, 1X10$^6$ cells were incubated in Fc receptor block (anti CD16) for 10 min and stained with CD45R, CD11c, CD11b, F4/80, Ly6C, Ly6G, Invitrogen Live dead stain (Aqua) in 1:200 ratio for 30 min in ice. Samples were washed thrice in FACS buffer and analyzed with a Cytek Aurora cytometer. Results were analyzed with FlowJo software (Tree Star, USA).

**Cytospin**

FACS sorted cells were centrifuged in 200 µL media onto a microscope slide using a Cytospin Universal 320 (Hettich, Germany) and stained with H&E stain. Images were acquired with a Nikon microscope equipped with a Canon A620 camera.

**Histology**

3h after DAP-muropeptide administration or 7 d after topical application of IMQ, pinnae were isolated and fixed in PBS containing 10% formalin. Paraffin-embedded sections were cut at 0.5 mm, stained with hematoxylin and eosin, and imaged by light microscopy.

**qRT-PCR**

Cells or tissue were lysed in TRIzol followed by RNA extraction per manufacture protocol. In brief, TRIzol lysed samples were mixed with chloroform and spun at 12000g for 15min at 4C. RNA from aqueous phase was precipitated with isopropanol and the RNA pellet was washed with 80% ethanol and resuspended in water.

cDNA was prepared from 1µg RNA using iScript gDNA Clear cDNA Synthesis Kit per manufacturer protocol. In brief, 1µg RNA was incubated with DNase mastermix at 25°C for 5 min and followed by reverse transcription for 20 min at 46C. cDNA was diluted in 1:5 and used directly in qPCR reaction.

Real-time quantitative PCR was performed with 0.4mM primer, 10 µL iQ SYBR Green Supermix (BioRAD), in a final volume of 10 µL on the CFX96 real-time system (BioRAD). All samples were run in triplicate. GAPDH was used to normalize. Primer sequences are reported in KRT.

**Isolation of RNA from mouse tissues and cells**

Lungs: whole lung was extracted from euthanized mice and 1mg lung tissue was homogenized in Trizol using homogenizer and cDNA was prepared as above.

Spleen: whole spleen was extracted from euthanized mice and single cell suspension was prepared by meshing through a 70 µm cell strainer with a plunger. These splenocytes were resuspended in Trizol and cDNA was prepared as above.

Gut: Gut was extracted from euthanized mice and 1 mg gut tissue was homogenized in Trizol using homogenizer and cDNA was prepared as above.

Dermis and epidermis: whole skin was extracted from euthanized mice and incubated 1mg/ml Dispase II (Roche) overnight at 4°C. Epidermis was separated from the dermis with forceps. These tissues were homogenized in Trizol and cDNA was prepared as above.

Bone marrow macrophages: tibia and femur bones were extracted from euthanized mice and crushed using mortar and pestle in DMEM medium. This suspension of cells was passed through a 70 µm cell strainer and $2 \times 10^6$ cells/mL were plated in DMEM F12 media with recombinant MCSF (25 ng/ml). After 7 days, cells were harvested in Trizol and cDNA was prepared as above.

Peritoneal macrophages: 2ml of 3% Brewer thioglycollate medium was injected to elicit peritoneal macrophages. 4 days later, peritoneal cells were harvested by injecting 5ml FACS buffer in peritoneal cavity of euthanized mice and collected. These cells were resuspended in Trizol and cDNA was prepared as above.

Dendritic cells: Pan dendritic cells were isolated from euthanized mice using Pan Dendritic Cell Isolation Kit from Miltenyi Biotec Inc. as per manufacturer protocol. cDNA was prepared as above.

Neutrophils: neutrophils were isolated from bone marrow cells. Marrow cells were isolated as described and neutrophils separated from them using histopaque density gradient (Histopaque 1119: Histopaque 1077 in 1:5(Sigma)). The middle layer containing neutrophils was separated and cells were resuspended in Trizol and cDNA was prepared as above.

**IMQ model of psoriasis-like dermatitis**

Psoriasis-like inflammation on mouse pinnae skin was induced as previously described (*12*). In brief, mice were treated daily for up to 6 d with 5 mg 5% IMQ cream topically applied on one pinna, while the other pinnae were treated with a similar amount of petroleum jelly, as vehicle control. Ear-skin thickness was measured daily using a digital caliper. The change in ear-skin thickness over time was reported as the difference related to measurement immediately prior to the first IMQ or VAS application. For bacterial association, $10^7$ CFU in 50% glycerol was applied daily to one pinna and 50% glycerol was applied to other pinna.

**Tape stripping**

The upper most layers of the epidermal barrier, the *stratum corneum,* of the dorsal pinnae was disrupted by tape stripping: manually applying a small piece of surgical tape (Transpore surgical tape) to the skin and removing, repeating 5 times with a fresh piece of tape for each round (*13*). Then$10^7$ cfu bacteria or 30µM iE-DAP in 20µl of 50% glycerol was applied to one pinna and only 50% glycerol was applied to another pinna. 3 h later, mice were euthanized, and pinnae were processed as described above.

**Click-iE DAP preparation and imaging**

iE-DAP and MDP with an alkyne handle for click-chemistry reaction were synthesized using a standard 8-step chemical synthesis, details to be published elsewhere.

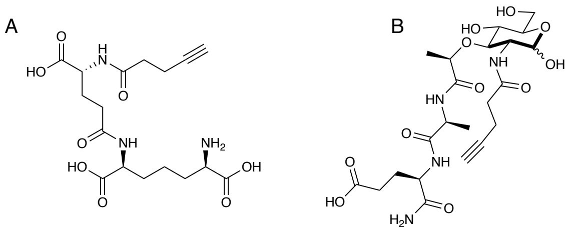


Structure of biologicaly active click-iE-DAP (A) and click-MDP (B): The probe compound contains an alkyne on the amino-terminus of the dipeptide and the 2-amino group of the muramic acid sugar.

The final click-muropeptides were purified by semi-prep C18 HPLC. Prior to use in experiments the final click-muropeptides were purified by semi-prep C18 HPLC and analyzed by NMR & MS. (Thermo Q-Exactive Orbitrap at the Mass Spectroscopy Facility and Bruker AV 400 MHz, AV III 600 MHz NMR at the NMR laboratory, Department of Chemistry and Biochemistry, University of Delaware.

When used to challenge primary keratinocytes, this click-iE DAP and -MDP was similarly active in inducing cell permeabilization compared to iE-DAP and MDP respectively (data not shown). For visualization, mouse keratinocytes were challenged with click-iE-DAP (30µM) or click-MDP (20µM) 37˚C for 30-60 min, cells washed 2X with 1xPBS to remove access of click-muropeptides and fixed with 4% paraformaldehyde in PBS at RT for 10 min. Cells were permeabilize with 1% Triton-X in PBS for 10 min at RT and blocked with 1% BSA in PBS. These permeabilized cells were then incubated in click-reaction conditions (250µM $CuSO_4$, 35µM BTAA, 60µM sodium ascorbate) with 2.5µM CalFluor 488 Azide at RT for 30 mins. Cells were then washed and mounted on slides, with DAPI containing mounting media. Slides were imaged with a Leica SP8 confocal microscope.

**Nano-particle preparation and challenge**

A lipid/cholesterol matrix consisting of 1,2-dioleoyl-sn-glycero-3-phosphocholine (DOPC, Avanti), 1,2-distearoyl-sn-glycero-3-phosphocholine (DSPC, Avanti), 1,2-dioleoyl-sn-glycero-3-phospho-(1’-rac-glycerol) (DOPG, Avanti), cholesterol (Avanti), and 1,2-distearoyl-sn-glycero-3-phosphoethanolamine-methoxyl poly(ethylene glycol) 2000 (DSPE-mPEG 2000, Laysan Bio) was used to formulate iE-DAP-encapsulated nanoparticles (NPs). We adapted our previously published methods (*14*) to load iE-DAP into lipid-based NPs. Briefly, a matrix of DOPC/DSPC/DOPG/cholesterol/DSPE-mPEG at 33.5/33.5/20/10/3 mol% was prepared in chloroform and dry lipid/cholesterol films were allowed to form. Films were rehydrated with iE-DAP (Invivogen) prepared in PBS at 1.2 mg/mL and these samples were vortexed for 30 sec every 10 min for 1 hr at 56ºC to complete this process. Following rehydration, samples were ultrasonicated in alternating 20 sec pulse/10 sec off cycles for 5 min at an amplitude of 20% to form iE-DAP-encapsulated lipid-based NPs. Samples were dialyzed for 1 hr following ultrasonication. Dynamic light scattering (DLS) and zeta potential measurements using a Malvern Zetasizer were used to characterize NP size and surface charge, respectively. NPs had an average 43.97 nm hydrodynamic diameter, a polydispersity index (PDI) of 0.158, and a zeta potential of -9.90 mV. Quant-IT assay (Thermo Fisher Scientific) measurements were used to measure average iE-DAP encapsulation at 998.1 µg/mL and an average encapsulation efficiency of 57.1%. To visualize iE-DAP-encapsulated NPs, 0.1 mol% of the fluorescent lipid tracer dye 3,3’-dioctadecyloxacarbocyanine perchlorate (DiO) were added to the lipid/cholesterol films.

Mouse keratinocytes were challenged iE-DAP loaded NPs, or control empty NPs, at 37˚C at 10μg/ml final concentration (~30 μM iE-DAP), and then monitored for Sytox uptake assay for 24 h, as described above. To visualize NP delivery, cells were washed after 30-60 mins 2X with PBS to remove access NPs and fixed with 4% paraformaldehyde in PBS at RT for 10 min. Cells were blocked with 1% BSA in PBS, washed 3X with PBS and mounted on slides with DAPI containing mounting media. Slides were then visualized with a Leica SP8 confocal microscope.

**Membrane permeability assays**
Mouse keratinocytes were plated at 5 x10ˆ4 cells/well of a tissue culture-treated 96 well plate with optically clear flat wells 24 hr prior to experimentation. Sytox and Hoechst 33342 were diluted in keratinocytes growth medium to a final concentration of 100 ng/ml and 0.3 μg/ml, respectively, and added to keratinocytes during treatment. For Sytox uptake cells were imaged using a Cytation 5 multi-mode reader from BioTek and analyzed by Gen5 software. Human keratinocytes were plated at 2.5-3 x10ˆ4 cells/well 24 hr prior to experimentation. Propidium iodide and Hoechst 33342 were diluted in KSFM to a final concentration of 1 μg/ml and 0.3 μg/ml, respectively, and added to keratinocytes during treatment. PI uptake was measured using a Cytation 5 multi-mode reader from BioTek and analyzed by Gen5 software.

**Human skin equivalent construction and infections**
HSEs were constructed as previously described (*15*). For muropeptides challenge assays, 30μM iE-DAP diluted in PBS was added to the epidermal side of the HSE. For IL-1R inhibition studies, 500 ng/ml of IL-1Ra (Biolegend) was added to 8 mL of HSE cornification media at 1 h post-iE-DAP challenge. Keratinocyte and fibroblast layers were separated by 5-10 min incubation in a dispase II containing buffer (150 mM NaCl, 10mM HEPES, 2mM $CaCl_2$, 2.4 U/ml dispase II) followed by gentle peeling of the two layers. Supernatants or individual tissues layers were harvested at 24 h for transcriptional analysis.

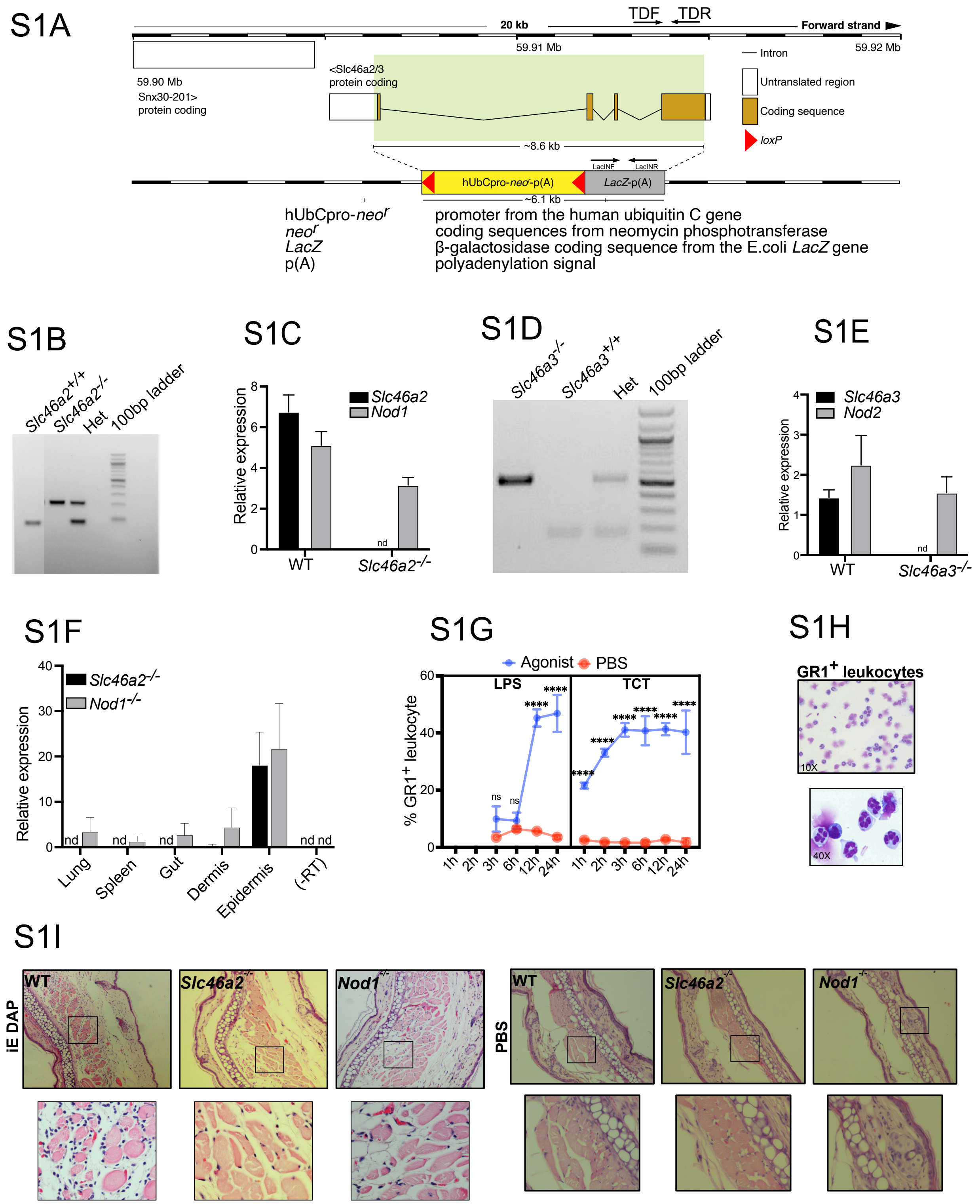


### Figure S1

**Fig. S1: *Slc46a2* Knockout Strategy and validation and recruitment of neutrophils in response to DAP muropeptide challenge.** (**A**) Design of *Slc46a2/3* null allele. The gene region was replaced with ZEN-UB1 targeting cassette by homologous recombination. Primers used for genotyping are indicated. Adapted from velocigene (www.velocigene.com) and Ensembl genome browser. (**B**) Agarose gel of PCR genotyping for validation of *Slc46a2* knockout mice. (**C**) Quantitative RT-PCR from WT and *Slc46a2*$^{-/-}$ mouse epidermis for *Slc46a2* and *Nod1* expression. *Slc46a2*$^{-/-}$ epidermis showed no *Scl46a2* expression. (**D**) Agarose gel of PCR genotyping for validation of *Slc46a3* knockout mice. (**E**) Quantitative RT-PCR from WT and *Slc46a3*$^{-/-}$ mouse epidermis for *Slc46a3* and *Nod1* expression. *Slc46a3*$^{-/-}$ epidermis showed no *Scl46a3* expression. (**F**) Expression analysis of *Slc46a2* and *Nod1* in indicated mouse organs. Maximum expression of both *Slc46a2* and *Nod1* was observed in epidermis. nd; not detected. (**G**) Neutrophil recruitment to the pinnae was measured at indicated time points after intradermal injection of10 µl of 10 µg/ml LPS or 8 µM TCT in one ear compared to a similar volume of PBS injection, as a control, in the contralateral ear. TCT triggered robust and rapid neutrophil recruitment in the skin while the response to LPS is slower. (**H**) Images of FACS sorted GR1$^{+}$ neutrophils 3 hr after iE-DAP challenge from WT mouse skin. Cells, prepared using cytospin and stained with Giemsa stain, show multilobed nuclei. (**I**) Representative H&E stained histological sections from the mouse ear skin after 3 hr intradermal injection of 10µl of 30µM iE-DAP or equal volume of PBS in WT, *Slc46a2*$^{-/-}$, and *Nod1*$^{-/-}$ mice. Inset shows the zoomed area of images. iE-DAP recruited inflammatory cells in WT but not in *Slc46a2*$^{-/-}$, and *Nod1*$^{-/-}$ mice skin, whereas PBS injection did not induce inflammatory reaction. Panel G uses two-way ANOVA and Tukey's multiple comparisons test to determine significance. Panels B-I are representative of at least three independent experimental results. **** $P < 0.0001$; ** $P < 0.01$; ns, not significant. $n \geq 3$ for all panels.

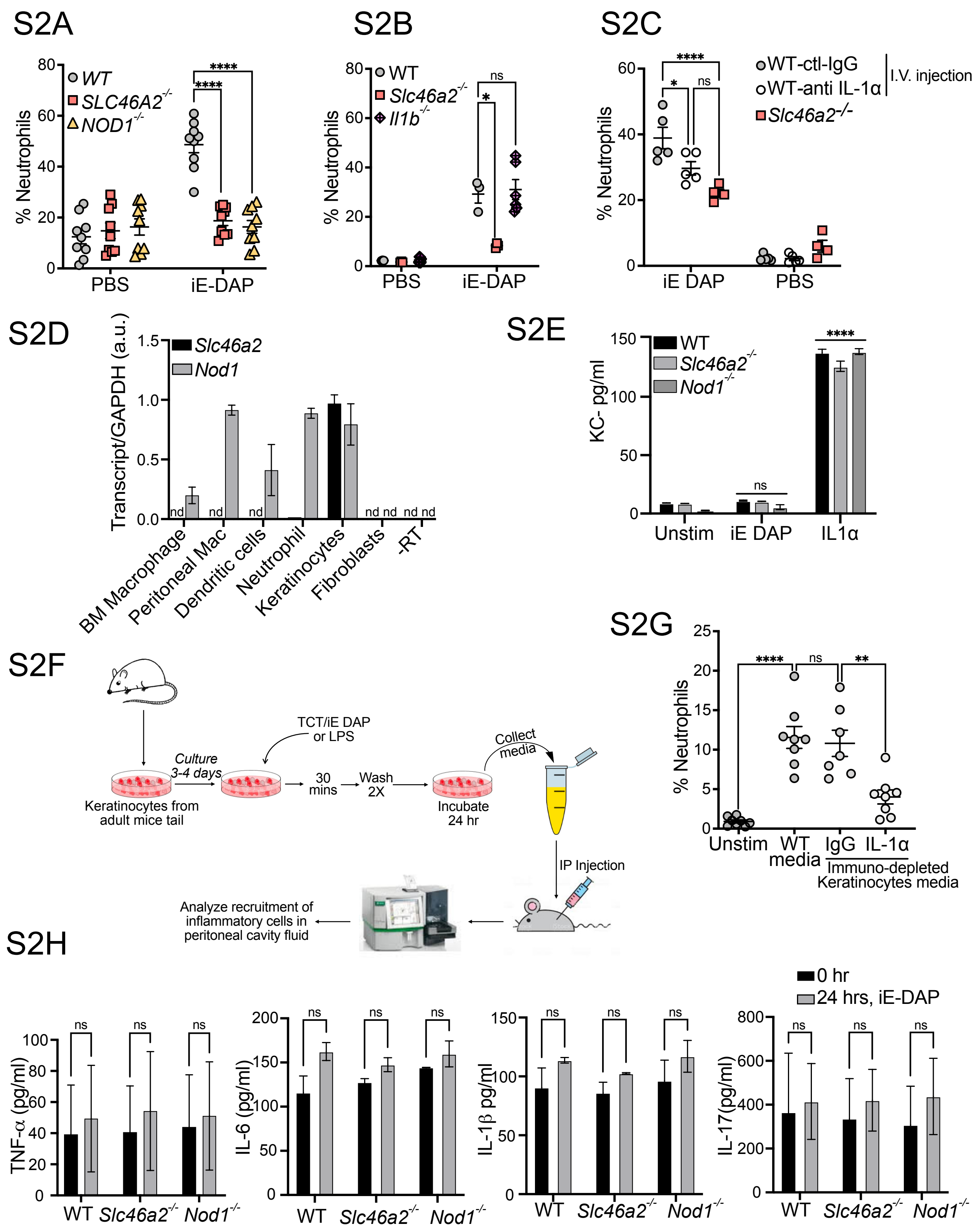


**Figure S2**

**Fig. S2: *Slc46a2*-dependent response to DAP-muropeptides recruits neutrophils and induces IL-1α.** (**A**) Neutrophil recruitment was measured in WT, *Slc46a2*$^{-/-}$ or

*Nod1*$^{-/-}$ mice pinnae 3 h after topical application of 10 µl of 30µM iE-DAP to tape stripped mouse skin. WT skin responded to iE-DAP challenge unlike *Slc46a2*$^{-/-}$ and *Nod1*$^{-/-}$ skin. (**B**) Neutrophil recruitment was measured in WT, *Il1b*$^{-/-}$ and *Slc46a2*$^{-/-}$ mice pinnae in response of intradermal challenge with 10 µl of 30µM iE-DAP. No significant difference was observed in GR1$^{+}$ cell recruitment in *IL1b*$^{-/-}$ mice compared to WT. (**C**) Mice were intravenously injected with IL-1α blocking antibody (1 µg/mouse), or isotype matched IgG control, 1 hour prior to intradermal injection of 10 µl of 30 µM iE-DAP, and 3 h later neutrophil recruitment was measured. Neutralization of IL-1α significantly reduced the recruitment of leukocytes in WT mice. (**D**) Expression analysis of *Slc46a2* and *Nod1* in specific cell types isolated from mice, by qRT-PCR. Highest expression of *Slc46a2* was observed in keratinocytes. (**E**) Primary dermal fibroblasts from WT, *Slc46a2*$^{-/-}$ and *Nod1*$^{-/-}$ mice were challenged with iE-DAP (30 µM) or IL-1α (10 ng/ml) for 24hrs and CXCL1 (KC) was measured by ELISA from culture media. KC was induced in fibroblasts treated with IL-1α, regardless of their *Slc46a2* or *Nod1* genotype, but fibroblasts were unresponsive to iE-DAP. **(F)** Schematic representation of experimental design for preparation of and bioassay analysis of conditioned media from primary mouse keratinocyte cultures. (**G**) Anti-IL-1α antibody was used to deplete this cytokine from keratinocyte conditioned media prior to IP injection. IL-1α depleted media showed reduced neutrophil recruitment compared to control IgG antibody-treated media. (**H**) TNFα, IL-6, IL-1β and IL-17 levels from WT, *Slc46a2*$^{-/-}$ and *Nod1*$^{-/-}$ keratinocyte media before and after challenge with 30 µM iE-DAP for 24hrs. No significant induction in any of these cytokines was detected. Panels A-C, E, G and H use two-way ANOVA and Tukey's multiple comparisons test to determine significance. Panels D, E, and H are representative of at least three independent experimental results. **** $P < 0.0001$; ** $P < 0.01$; ns, not significant. $n \geq 3$ for all panels.

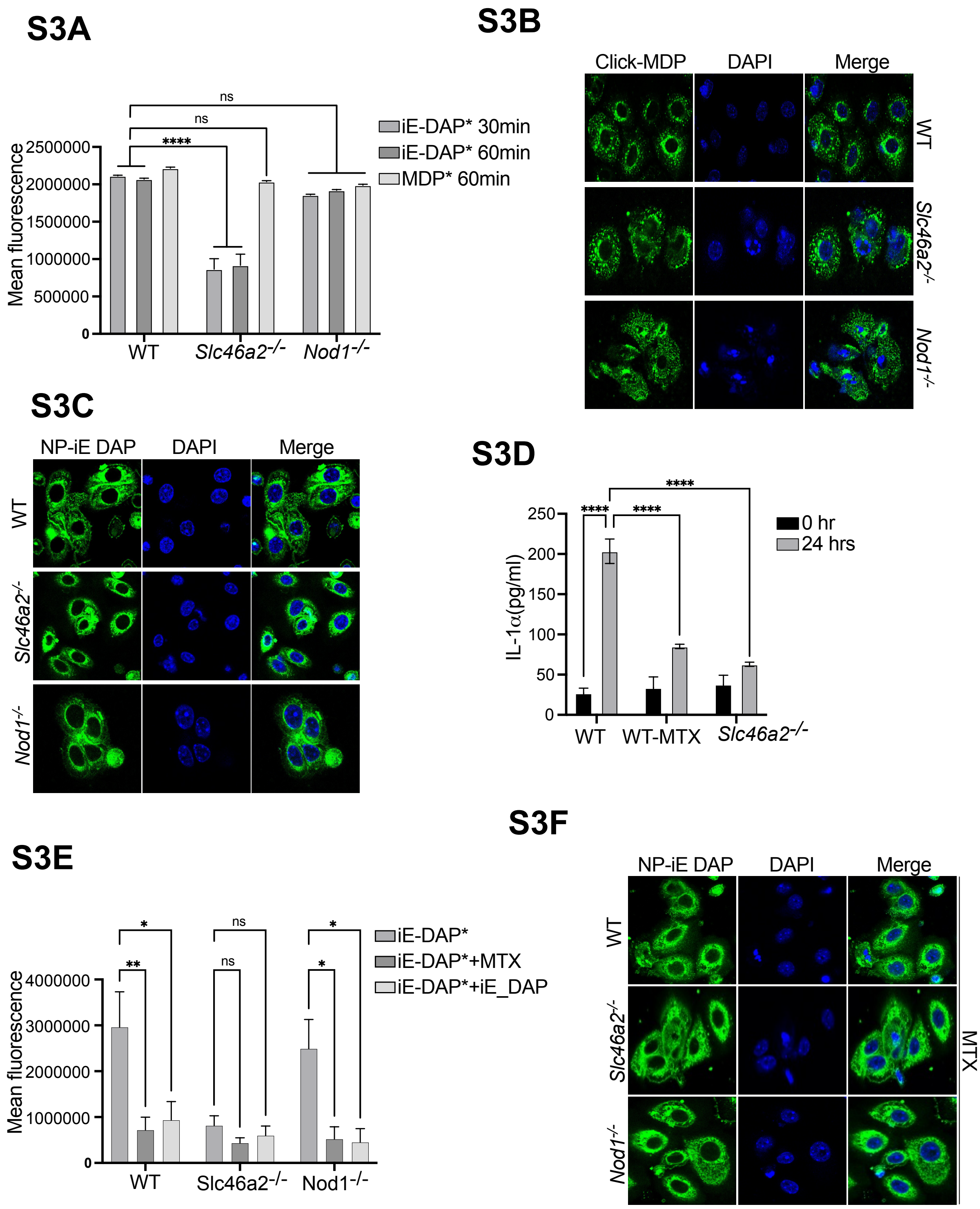


#### Figure S3

**Fig. S3: MTX blocks the transport of DAP muropeptides through SLC46A2.** (**A**) WT, *Slc46a2*$^{-/-}$ and *Nod1*$^{-/-}$, keratinocytes were challenged with 30 µM click-iE DAP or

10μM click-MDP for 30 minutes and 60 minutes, washed extensively, and then lysed. Muropeptides were then detected in these lysates with click-reacted CalFluor 488 Azide and fluorescence quantified. With iE-DAP, *Slc46a2*$^{-/-}$ keratinocytes showed significantly reduced fluorescence intensity compared to WT or *Nod1*$^{-/-}$ keratinocytes, while no change in fluorescence intensity was observed with click-MDP in all genotypes. (**B**) Fluorescent confocal microscopy of WT, *Slc46a2*$^{-/-}$ and *Nod1*$^{-/-}$ keratinocytes, challenged with 10μM "click-MDP" for 1h. WT, *Slc46a2*$^{-/-}$ and *Nod1*$^{-/-}$keratinocytes shows similar import of click-MDP. (**C**) Fluorescent confocal microscopy of WT, *Slc46a2*$^{-/-}$ and *Nod1*$^{-/-}$ primary keratinocytes treated with NPs loaded with iE-DAP and immunofluorescence dye. Localization of dye inside the keratinocytes shows the successful delivery NP delivery of cargo. No change in fluorescence intensity was observed in all genotypes. (**D**) IL-1α in culture media of WT and *Slc46a2*$^{-/-}$ keratinocytes after stimulating with 30 μM iE DAP for 24 hours. WT keratinocytes were also treated with 250 μM MTX along with iE-DAP. Like *Slc46a2*-deficiency, MTX prevented IL-1α release in iE-DAP treated WT keratinocytes. (**E**) Similar to (A), MTX (250μM) or unlabeled iE-DAP (30μM) interfered with the cellular uptake of click-iE-DAP in WT or *Nod1*$^{-/-}$, keratinocytes, while in *Slc46a2*$^{-/-}$ cells import was low and unchanged. (**F**) Immunofluorescent images of WT, *Slc46a2*$^{-/-}$ and *Nod1*$^{-/-}$ keratinocytes pretreated with 250 μM MTX and then iE-DAP delivered by dye-loaded NP. Addition of MTX did not affect the NP mediated dye delivery in any genotype. Panels A, D and E use two-way ANOVA and Tukey's multiple comparisons test to determine significance. Panels B, C and F are representative of at least three independent experimental results. **** $P < 0.0001$; ** $P < 0.01$; ns, not significant. $n \geq 3$ for all panels.

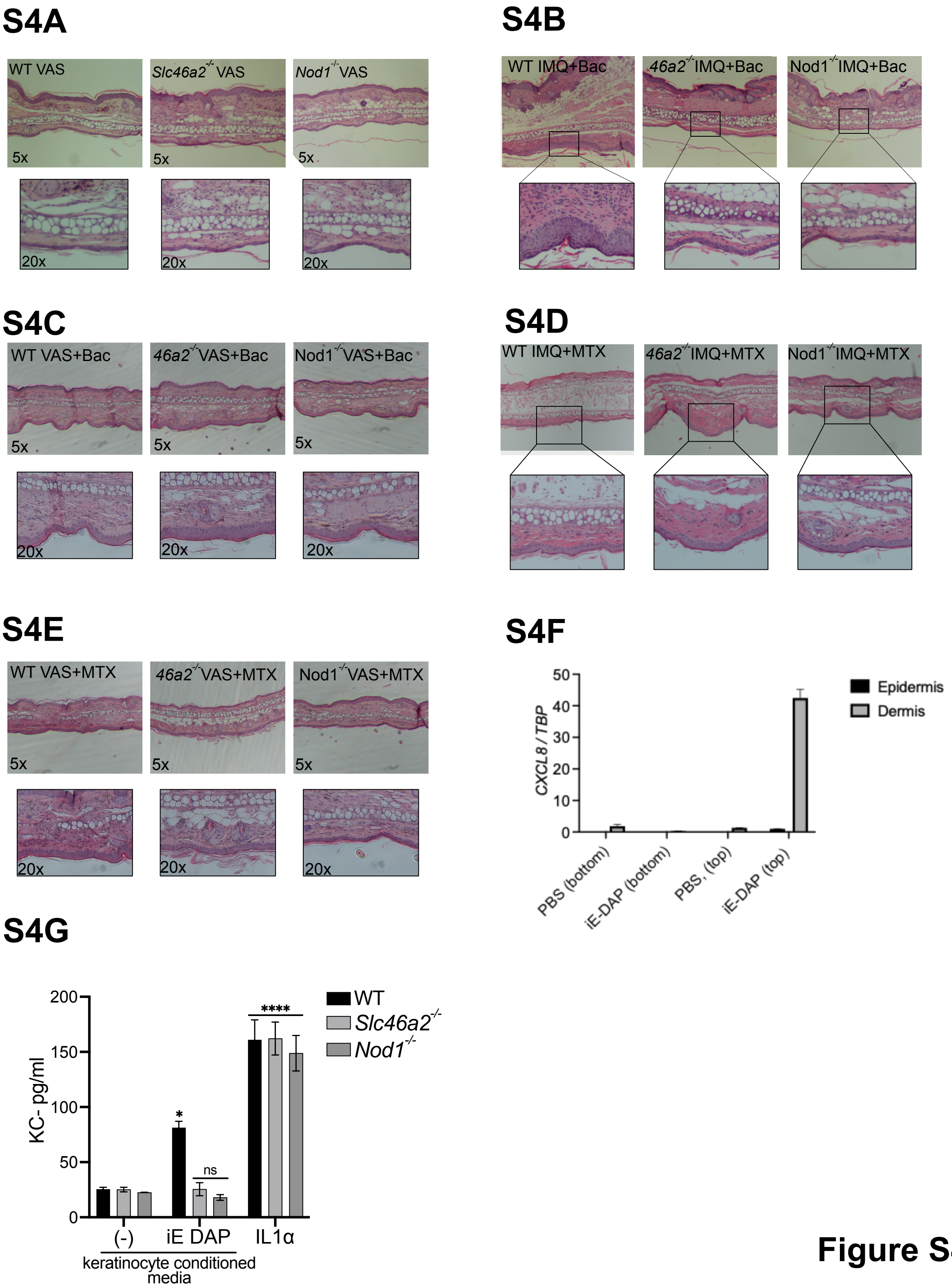


**Figure S4**

**Fig. S4: MTX blocks the psoriatic inflammation in skin**. (**A**) Representative H&E stained histology of ear sections from VAS (Vaseline, as vehicle) applied skin from WT, *Slc46a2*$^{-/-}$ and *Nod1*$^{-/-}$ mice. VAS applied skin did not show signs of inflammation in any genotype. (**B**) H&E stained histology sections from IMQ and *C. accolens* (Bac) applied skin. WT skin shows hyper inflammation compared to *Slc46a2*$^{-/-}$ and *Nod1*$^{-/-}$ mice skin. (**C**) Representative H&E stained histology of ear sections from VAS (Vaseline, as vehicle) and *C. accolens* (Bac) applied skin from WT, *Slc46a2*$^{-/-}$ and *Nod1*$^{-/-}$ mice. VAS applied skin did not show signs of inflammation in any genotype. (**D**) H&E stained histology sections from IMQ and methotrexate (MTX) treated skin from from WT, *Slc46a2*$^{-/-}$ and *Nod1*$^{-/-}$ mice. After application of MTX WT skin was less inflamed. However, *Slc46a2*$^{-/-}$ and *Nod1*$^{-/-}$ skin has a minimalistic effect of MTX. (**E**) Representative H&E stained histology of ear sections from VAS (Vaseline, as vehicle) and methotrexate (MTX) applied skin from WT, *Slc46a2*$^{-/-}$ and *Nod1*$^{-/-}$ mice. VAS applied skin did not show signs of inflammation in any genotype. (**F**) CXCL8 expression was analyzed in skin organoids after iE-DAP challenge in the top (epidermal layer) or bottom (dermal layer). CXCL8 response was only observed in dermal fibroblast when an iE-DAP challenge was given to the epidermal keratinocytes. (**G**) WT primary dermal fibroblasts were cultured for 24 h in conditioned media from WT, *Slc46a2*$^{-/-}$ or *Nod1*$^{-/-}$ keratinocytes, that were stimulated or not with 30 μM iE-DAP. As a control WT, *Slc46a2*$^{-/-}$ and *Nod1*$^{-/-}$ fibroblasts were treated with IL1-α (10ng/ml). KC was induced in fibroblasts cultured in condition media from WT keratinocytes treated with iE-DAP or fibroblasts treated with IL1-α. Panel G uses two-way ANOVA and Tukey's multiple comparisons test to determine significance. Panels A-F are representative of at least three independent experimental results. **** $P < 0.0001$; ** $P < 0.01$; ns, not significant. $n \geq 3$ for all panels.

#### Bibliography for Supplemental Methods


1. D. M. Valenzuela *et al.*, High-throughput engineering of the mouse genome coupled with high-resolution expression analysis. *Nat Biotechnol* **21**, 652-659 (2003).
2. K. S. Kobayashi *et al.*, Nod2-dependent regulation of innate and adaptive immunity in the intestinal tract. *Science* **307**, 731-734 (2005).
3. M. Chamaillard *et al.*, An essential role for NOD1 in host recognition of bacterial peptidoglycan containing diaminopimelic acid. *Nat Immunol* **4**, 702-707 (2003).
4. N. Kayagaki *et al.*, Caspase-11 cleaves gasdermin D for non-canonical inflammasome signalling. *Nature* **526**, 666-671 (2015).
5. T. Kawai, O. Adachi, T. Ogawa, K. Takeda, S. Akira, Unresponsiveness of MyD88-deficient mice to endotoxin. *Immunity* **11**, 115-122 (1999).
6. S. M. Man *et al.*, Differential roles of caspase-1 and caspase-11 in infection and inflammation. *Sci Rep* **7**, 45126 (2017).
7. M. B. Glaccum *et al.*, Phenotypic and functional characterization of mice that lack the type I receptor for IL-1. *J Immunol* **159**, 3364-3371 (1997).
8. V. A. K. Rathinam *et al.*, The AIM2 inflammasome is essential for host defense against cytosolic bacteria and DNA viruses. *Nature Immunology* **11**, 395-402 (2010).
9. R. K. S. Malireddi *et al.*, Determining distinct roles of IL-1α through generation of an IL-1α knockout mouse with no defect in IL-1β expression. *bioRxiv*, 2022.2009.2021.508892 (2022).
10. F. Li, C. A. Adase, L. J. Zhang, Isolation and Culture of Primary Mouse Keratinocytes from Neonatal and Adult Mouse Skin. *J Vis Exp*, (2017).
11. A. Ray, B. N. Dittel, Isolation of mouse peritoneal cavity cells. *J Vis Exp*, (2010).
12. E. E. Gray *et al.*, Deficiency in IL-17-committed Vgamma4(+) gammadelta T cells in a spontaneous Sox13-mutant CD45.1(+) congenic mouse substrain provides protection from dermatitis. *Nat Immunol* **14**, 584-592 (2013).
13. F. Au - Sparber, S. Au - LeibundGut-Landmann, Infecting Mice with Malassezia spp. to Study the Fungus-Host Interaction. *JoVE*, e60175 (2019).
14. P. U. Atukorale *et al.*, Nanoparticle Encapsulation of Synergistic Immune Agonists Enables Systemic Codelivery to Tumor Sites and IFNbeta-Driven Antitumor Immunity. *Cancer Res* **79**, 5394-5406 (2019).
15. M. W. Carlson, A. Alt-Holland, C. Egles, J. A. Garlick, Three-dimensional tissue models of normal and diseased skin. *Current Protocols in Cell Biology* **41**, 19.19. 11-19.19. 17 (2008).
